# Secretory leukocyte protease inhibitor influences periarticular joint inflammation in *B. burgdorferi*-infected mice

**DOI:** 10.1101/2024.11.24.625079

**Authors:** Qian Yu, Xiaotian Tang, Thomas Hart, Robert Homer, Alexia A. Belperron, Linda K. Bockenstedt, Aaron Ring, Akira Nakamura, Erol Fikrig

**Affiliations:** Section of Infectious Diseases, Department of Internal Medicine, School of Medicine, Yale University, New Haven, Connecticut, USA; Department of Pathology, School of Medicine, Yale University, New Haven, Connecticut, USA; Section of Rheumatology, Allergy and Immunology, Department of Internal Medicine, School of Medicine, Yale University, New Haven, Connecticut, USA; Department of Immunobiology, Yale School of Medicine, New Haven, CT, USA; Department of Pharmacology, Yale School of Medicine, New Haven, CT, USA; Divisions of Immunology, Faculty of Medicine, Tohoku Medical and Pharmaceutical University, Sendai, Japan; Institute of Insect Sciences, College of Agriculture and Biotechnology, Zhejiang University, Hangzhou, China

## Abstract

Lyme disease, caused by *Borrelia burgdorferi*, is the most common tick-borne infection in the United States. Arthritis is a major clinical manifestation of infection, and synovial tissue damage has been attributed to the excessive pro-inflammatory responses. The secretory leukocyte protease inhibitor (SLPI) promotes tissue repair and exerts anti-inflammatory effects. The role of SLPI in the development of Lyme arthritis in C57BL/6 mice, which can be infected with *B. burgdorferi*, but only develop mild joint inflammation, was therefore examined. *SLPI*-deficient C57BL/6 mice challenged with *B. burgdorferi* had a higher infection load in the tibiotarsal joints and marked periarticular swelling, compared to infected wild type control mice. The ankle joint tissues of *B. burgdorferi-*infected *SLPI*-deficient mice contained significantly higher percentages of infiltrating neutrophils and macrophages. *B. burgdorferi*-infected *SLPI*-deficient mice also exhibited elevated serum levels of IL-6, neutrophil elastase, and MMP-8. Moreover, using a recently developed BASEHIT (**BA**cterial **S**election to **E**lucidate **H**ost-microbe **I**nteractions in high **T**hroughput) library, we found that SLPI directly interacts with *B. burgdorferi*. These data demonstrate the importance of SLPI in suppressing periarticular joint inflammation in Lyme disease.

## Introduction

Lyme disease is the most common tick-borne illness in the United States, affecting an estimated 500,000 people each year^1^. The spirochete *Borrelia burgdorferi* is the causative agent of Lyme disease and is primarily transmitted by *Ixodes scapularis* ticks in North America^2^. Early administration of antibiotics is usually successful in the treatment of Lyme disease. However, between 2008-2015, arthritis was the major manifestation in a third of Lyme disease cases reported to CDC^3,4^. Musculoskeletal symptoms occur at all stages of Lyme disease, with migratory arthralgias in the early stages and frank arthritis occurring months later. Lyme arthritis can present as acute or intermittent self-resolving episodes or persistent joint swelling and pain, which if left untreated, can lead to irreversible joint dysfunction and debilitation^3,5,6^. Although Lyme arthritis resolves completely with antibiotic therapy in most patients, a small percentage of individuals experience persistent joint inflammation for months or several years, termed post-infectious Lyme arthritis^3,5,7^.

Studies of synovial fluid from Lyme arthritis patients found infiltrating polymorphonuclear cells (PMNs), IFN-γ-producing mononuclear cells, and large amounts of NF-κB-induced pro-inflammatory cytokines and chemokines, such as IL-6, CXCL10, and TNF-α^6,8–10^. An inverse correlation between the robust IFN-γ signature and tissue repair has been demonstrated in the synovial tissue and fluid from patients with post-infectious Lyme arthritis^11^. This suggests that the dysregulated excessive pro-inflammatory responses inhibit tissue repair and lead to extensive tissue damage.

*B. burgdorferi* infection of laboratory mice causes an acute arthritis, the severity of which is mouse strain dependent^12^. *B. burgdorferi* infected-C3H/HeN mice develop pronounced neutrophilic infiltration of periarticular structures and the synovial lining which peaks in severity several weeks after infection^13^. In contrast, infection of *B. burgdorferi* C57BL/6 mice causes mild, if any, arthritis^14^. On a C57BL/6 background, the immune-deficient RAG−/− and SCID mice are also resistant to *B. burgdorferi*-induced arthritis, indicating that responses independent of humoral and cellular immunity contribute to the milder phenotype of disease in these animals^15^. Similar to Lyme arthritis in humans, neutrophils, macrophages, and signaling involving IFN-γ and NF-κB contribute to the severity of murine joint inflammation^16–19^.

The secretory leukocyte protease inhibitor (SLPI) is a 12 kDa, secreted, non-glycosylated, cysteine-rich protein^20^. It strongly inhibits serine proteases, especially neutrophil-derived serine proteases (NSPs) including cathepsin G (CTSG) and elastase (NE)^21,22^. It is secreted by epithelial cells at various mucosal surfaces and is also produced by neutrophils, macrophages, mast cells, and fibroblasts^23,24^. The major function of SLPI is to inhibit excessive protease activity at sites of inflammation, thus promoting tissue repair and wound healing^25,26^. SLPI also exerts anti-inflammatory function by inhibiting NF-κB activation in macrophages^27–29^. The roles of neutrophils and NSPs have been extensively studied in rheumatoid arthritis, a condition sharing some similarities with Lyme arthritis^30,31^. NE and CTSG induce potent destruction of cartilage proteoglycan *in vitro* and *in vivo*, which contributes to rheumatoid arthritis progression^32^. Some studies also demonstrated that SLPI inhibits joint inflammation and bone destruction^33,34^. However, the importance of SLPI and NSPs have not been studied in the context of Lyme disease.

Thus, in this study, we examined the role of SLPI in the development of murine Lyme arthritis caused by *B. burgdorferi*. Using the *SLPI*-deficient C57BL/6 mice, we observed a significant increase in the infection burden and marked periarticular swelling in the ankle joints compared to WT mice following *B. burgdorferi* infection. Significant increase in infiltrating neutrophils and macrophages were observed in the ankle joints of infected *SLPI*-deficient mice. Elevated serum levels of IL-6, neutrophil elastase, and MMP-8 in the infected *SLPI*-deficient mice were also observed, which can lead to the recruitment of neutrophils and macrophages exacerbating the periarticular swelling. We further demonstrated the direct interaction between SLPI and *B. burgdorferi*. This is the first study showing the importance of anti-protease-protease balance in the development of murine Lyme arthritis.

## Results

### Secretory leukocyte protease inhibitor (SLPI) influences periarticular joint inflammation in B. burgdorferi-infected mice

To assess the importance of SLPI during murine Lyme arthritis, we compared the outcomes of *B. burgdorfe*ri infection of C57BL/6 WT and *SLPI*−/− mice. The C57BL/6 WT and *SLPI*−/− mice were infected with 10^5^ spirochetes subcutaneously. Infection burdens in the skin were assessed by qPCR of *B. burgdorferi* DNA in ear punch biopsies at 7, 14, and between 21 to 24 dpi (Figure 1, A-C). Infection burdens in the heart (Figure 1D) and tibiotarsal joint (Figure 1E) tissues were assessed between 21 to 24 dpi. We did not observe any significant difference in infection burden in the skin between WT and *SLPI*−/− mice (n=24) at 7, 14, 21 to 24 dpi, or in the heart between 21 to 24 dpi (Figure 1, A-D).

**Figure 1.**
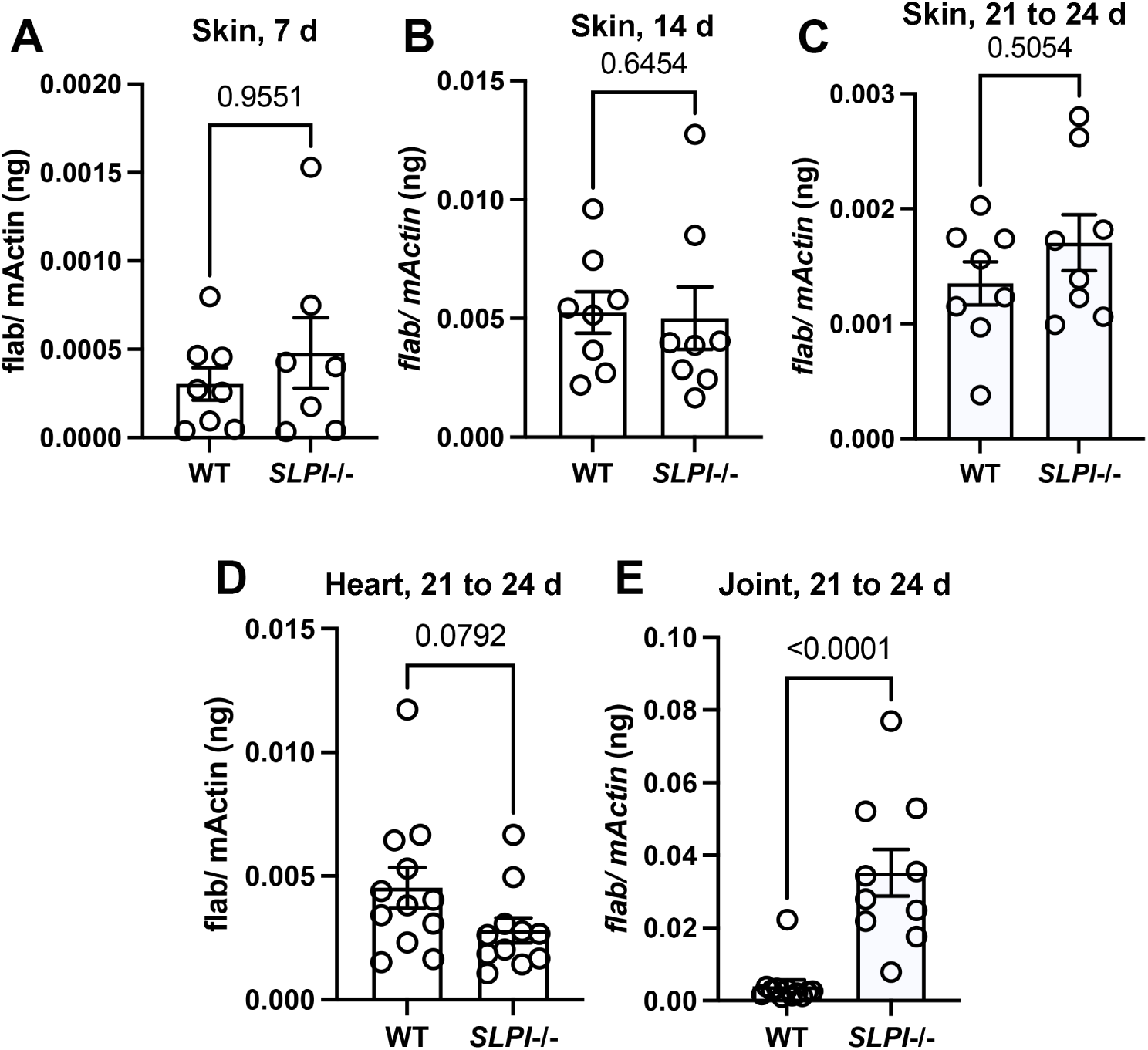
*B. burgdorferi* burden in C57BL/6 WT and *SLPI*−/− mice. WT and *SLPI*−/− mice were infected with 10^5^ spirochetes by subcutaneous injection. (A-C) Spirochete burden in skin was assessed by ear punch biopsies at 7d (A) 14 d (B), and between 21 to 24 d (C) post infection. (D and E) Spirochete burden in tibiotarsal joint and heart tissues was assessed between 21 to 24 d (D, heart, E, joint) post infection. At least n = 6 mice were infected in each group. The spirochete burden was measured by qPCR detecting *flaB* and normalized to mouse β*-actin*. Each data point represents an individual animal. Representative data were shown from three separate experiments. The error bars represent mean ± SEM and *p*-values are calculated using the non-parametric Mann-Whitney test.

Strikingly, we observed a significantly higher spirochete burden in the ankle joints of infected *SLPI*−/− mice (n=24) between 21 to 24 dpi (Figure 1E). Furthermore, at around 24 dpi, significant swelling was also observed solely in the infected *SLPI*−/− mice (Figure 2A, red arrow). The level of swelling was first scored visually. While 14 out of 20 infected *SLPI*−/− mice displayed visible swelling (score ≥ 1), only 1 in 14 infected WT mice displayed mild swelling (score 1) at the ankle (Figure 2B). The tibiotarsal joints were then dissected, fixed, and stained with H&E for histopathological evaluation of the level of inflammation (Figure 2, C and D). Inflammation of bursa and soft tissue adjacent to the tibiotarsal joint, but not in the tibiotarsal synovium, was consistently observed in the infected *SLPI*−/− mice (Figure 2C, black rectangle). In contrast, only 1 out of 14 infected WT mice displayed modest inflammation (score = 2) in these sites (Figure 2D). The above data indicate the importance of SLPI in modulating the development of periarticular inflammation associated with murine Lyme arthritis.

**Figure 2.**
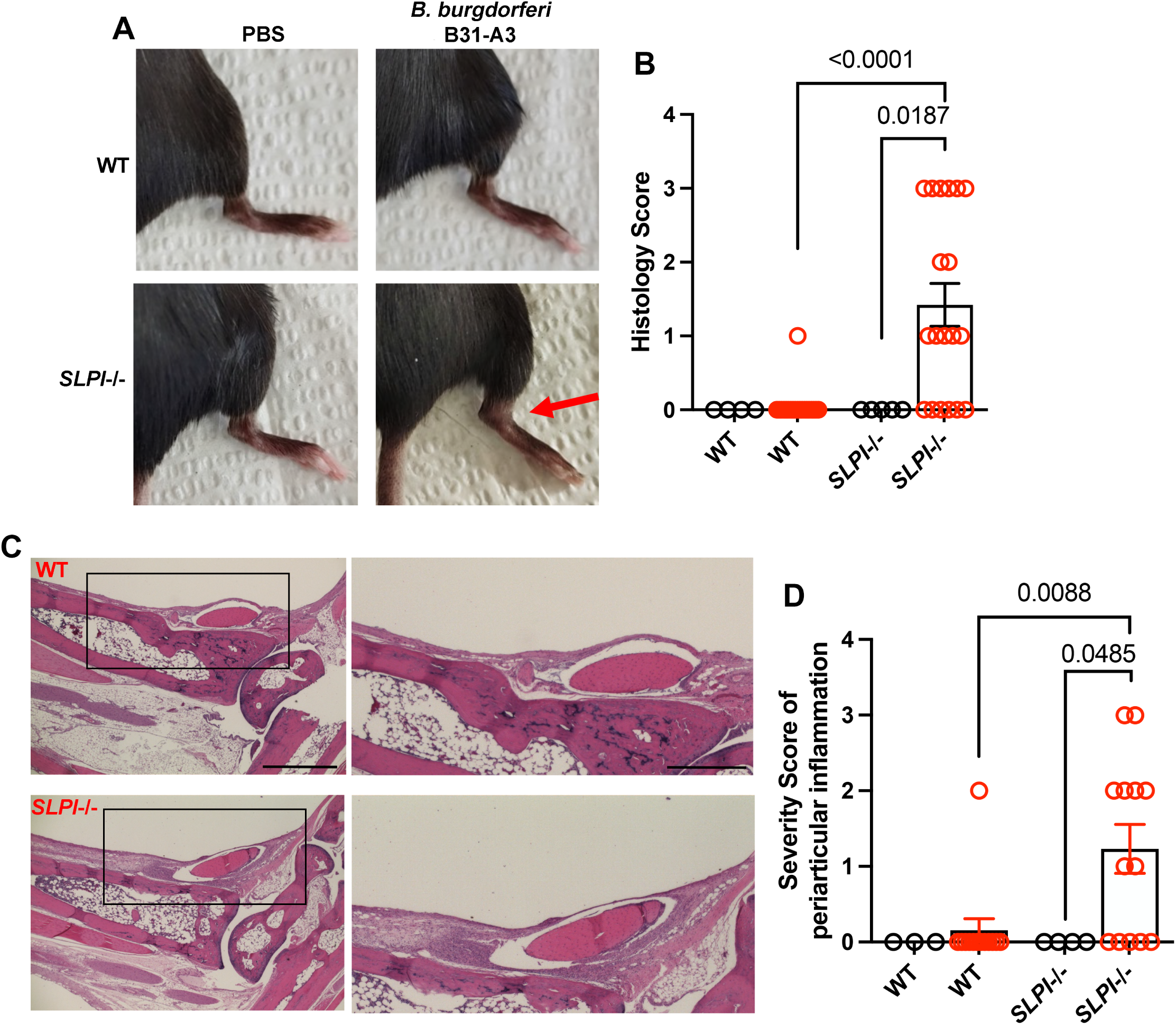
Assessment of ankle inflammation in WT and *SLPI*−/− mice infected with *B. burgdorferi* between 21 to 24 dpi. (A) Representative images are shown of the tibiotarsal joints of WT and *SLPI*−/− mice with/without *B. burgdorferi* infection between 21 to 24 dpi. The swelling is indicated by the red arrow. (B) Swelling of the tibiotarsal joints of individual mice was scored visually by an observer blinded to the experimental groups (scale of 0 (negative) to 3 (severe)). (C) The tibiotarsal joint of each mouse was dissected, fixed, sectioned, and stained with H&E. Representative images from *B. burgdorferi*-infected C57BL/6 WT and *SLPI*−/− mice are shown. Lower magnification (left panels, Scale bar: 100 μm) and higher magnification (right panels, Scale bar: 50 μm) of selected areas (black rectangle) are shown. (D) The severity of periarticular inflammation was scored blindly by the pathologist on a scale of 0 (negative) to 3 (severe). black, PBS-sham infection; red, *B. burgdorferi* infection. Results from two independent experiments were pooled and shown here. The error bars represent mean ± SEM and *p*-values are calculated using the non-parametric Mann-Whitney test.

### SLPI influences immune responses in B. burgdorferi-infected mice

It has been established that SLPI exerts its anti-inflammatory effect by inhibiting neutrophil serine protease and by dampening NF-κB activation in macrophages^22^. Thus, to investigate the mechanism underlying the effect of SLPI on murine joint inflammation, we sought to identify the population of infiltrating cells in the periarticular tissues of infected WT and *SLPI*−/− mice. Between 21 to 24 dpi, the ankle joints were dissected. To obtain single cell suspensions of infiltrating cells, bone marrow cells were removed and discarded and the joint and periarticular tissues were digested^35^. The cells were stained for flow cytometry with CD45, CD11b, and LY6G to label neutrophils (Figure 3A), and CX3CR1, CD64, and LY6C to label macrophages (Figure 3B)^36^. After gating, we observed significantly higher percentages of infiltrating neutrophils and macrophages in the dissected tissues from infected *SLPI*−/− than WT mice (Figure 3, A and B). To further eliminate the possibility of neutrophilic contamination within the macrophage population, we also implemented a Ly6G-negative gating strategy. The result showed a consistently higher percentage of macrophages in the infected *SLPI*−/− mice (Supplemental Figure 1). Using RT-qPCR on the tibiotarsal tissues extracted from *B. burgdorferi*-infected *SLPI*−/− mice, we detected increased gene expression of neutrophil chemoattractant receptor C-X-C motif chemokine receptor 2 (*CXCR2)*, monocyte chemoattractant protein 1 (*MCP-1*), and its receptor C-C motif chemokine receptor 2 (*CCR2)* (Figure 3, C-E).

**Figure 3.**
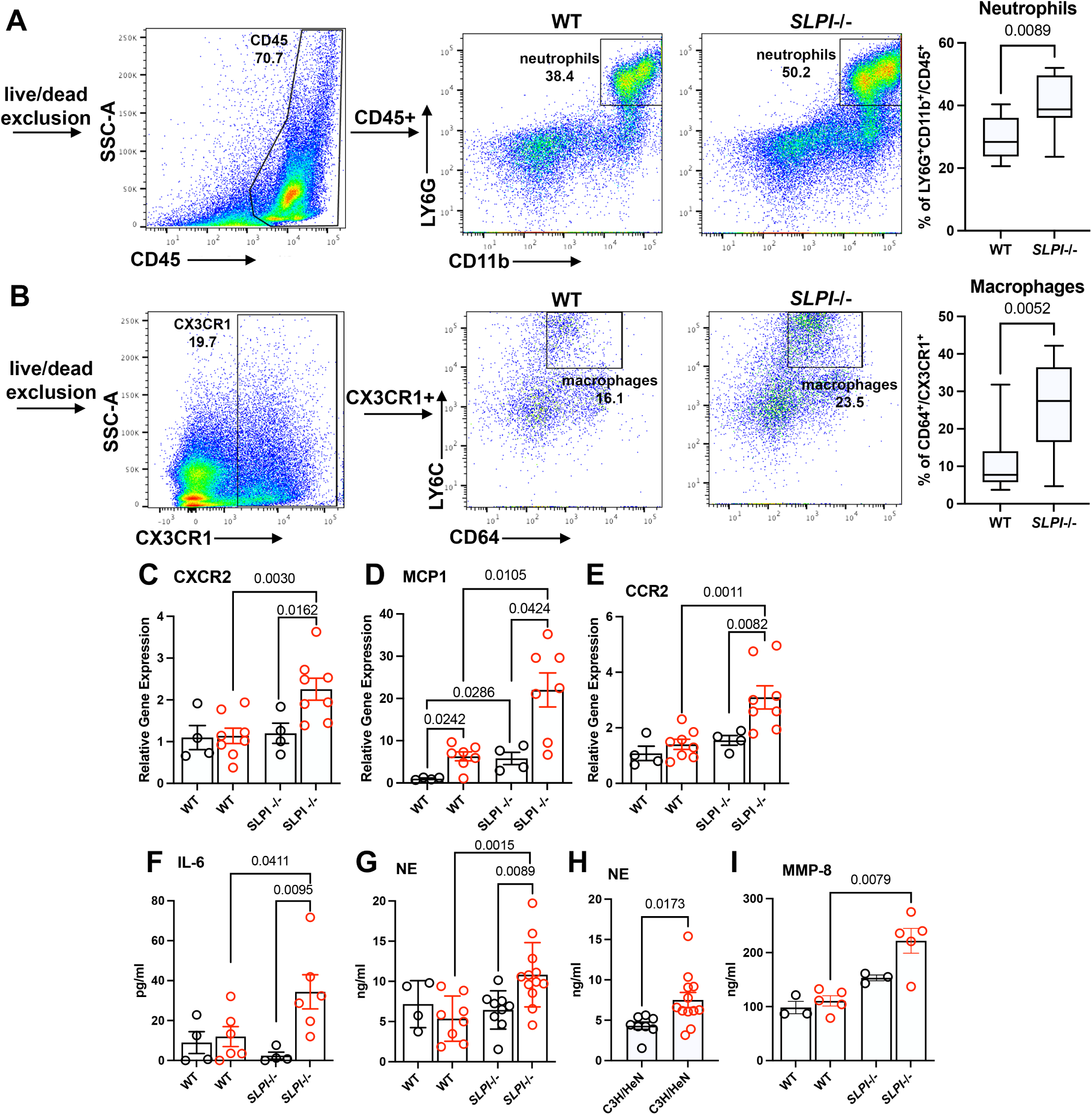
Immune profile analysis of infected WT and *SLPI*−/− mice. (A, B) Infiltrating cell population analysis of tibiotarsal joint tissues of infected WT and *SLPI*−/− mice. (A) The neutrophil population was gated on the CD11bLY6G double positive cells among the CD45 positive cells. (B) The macrophage population was gated on the CD64 positive cells among the CX3CR1 positive myeloid cells. Results from two independent experiments were pooled and shown here. (C-E) Expression levels of C-X-C motif chemokine receptor 2 (*CXCR2*, C), monocyte chemoattractant protein 1 (*MCP-1*, D), and C-C motif chemokine receptor 2 (*CCR2*, E) were assessed in the tibiotarsal tissue using RT-qPCR. (F) The serum cytokine profile was assessed using mouse cytokine/chemokine 32-plex array. An increase in IL-6 was observed in the infected *SLPI*−/− mice. (G, H) The serum level of neutrophil elastase (NE) was measured using an ELISA kit. (I) Serum levels of MMPs were assessed using a mouse MMP 5-Plex Discovery Assay. An increase in MMP-8 was observed in the infected *SLPI*−/− mice. Serum was obtained by cardiac puncture of WT and *SLPI*−/− C57BL/6 mice with/without infection between 21 to 24 dpi (F, G, and I) and of infected C3H/HeN mice at 21 dpi (H). black, PBS-sham infection; red, *B. burgdorferi* infection. Each data point represents an individual animal. The error bar represents mean ± SEM and *p*-values are calculated using the non-parametric Mann-Whitney test.

Furthermore, the serum cytokine/chemokine profile was assessed from uninfected and *B. burgdorferi*-infected WT and *SLPI*−/− mice between 21 to 24 dpi. We observed a significant increase in IL-6 in infected *SLPI*−/− mice (Figure 3F). IL-6 recruits and stimulates neutrophils, leading to secretion of neutrophil-derived serine proteases including neutrophil elastase (NE) and cathepsin G (CTSG)^37^. The lack of serine protease inhibitors, such as SLPI, can cause excessive protease activity and subsequent tissue damage and inflammation^31^. Indeed, using ELISA, we observed a significantly higher level of NE solely in the serum of infected *SLPI*−/− mice (Figure 3G). An increased serum level of NE was also observed in the *B. burgdorferi*-infected, arthritis-susceptible C3H/HeN mice at 21 dpi (Figure 3H). These data suggest that, in the absence of SLPI, excessive serine protease activity can exacerbate murine Lyme arthritis.

A correlation between IL-6, macrophages, metalloproteinases (MMPs), and articular cartilage destruction has been observed in the synovial tissue of RA patients^38^. Elevated levels of host matrix metalloproteinases (MMPs) have also been found in the synovial fluid of Lyme arthritis patients and can cause excessive tissue damage^39^. Thus, a mouse MMP 5-Plex Discovery Assay was used to explore the serum levels of different MMPs. We observed a significant increase in the levels of MMP-8 solely in the infected *SLPI*−/− mice (Figure 3I). Taken together, our data suggest that SLPI suppresses inflammation in *B. burgdorferi*-infected mouse joint tissues by potentially inhibiting neutrophil and macrophage infiltration and subsequent protease-mediated tissue destruction.

### Decreased serum level of SLPI in Lyme disease patients

Despite numerous studies of serum, synovial fluid and tissue from Lyme arthritis patients, the importance of anti-protease-protease balance in Lyme arthritis has not been investigated^7^. Based on our data obtained from the *SLPI*-deficient mice, we assessed the serum SLPI level in Lyme disease patients (Figure 4). Due to the limited samples available from Lyme arthritis patients, we included samples from Lyme disease patients who presented with earlier manifestations of Lyme disease. The serum level of human SLPI assessed by ELISA showed a significant decrease in the SLPI level in Lyme disease patients comparing to healthy adult controls (Figure 4). Similar to our data from *B. burgdorferi*-infected mice, this result suggests a correlation between the lack of SLPI and humans exhibiting clinical manifestations of Lyme disease, including arthritis.

**Figure 4.**
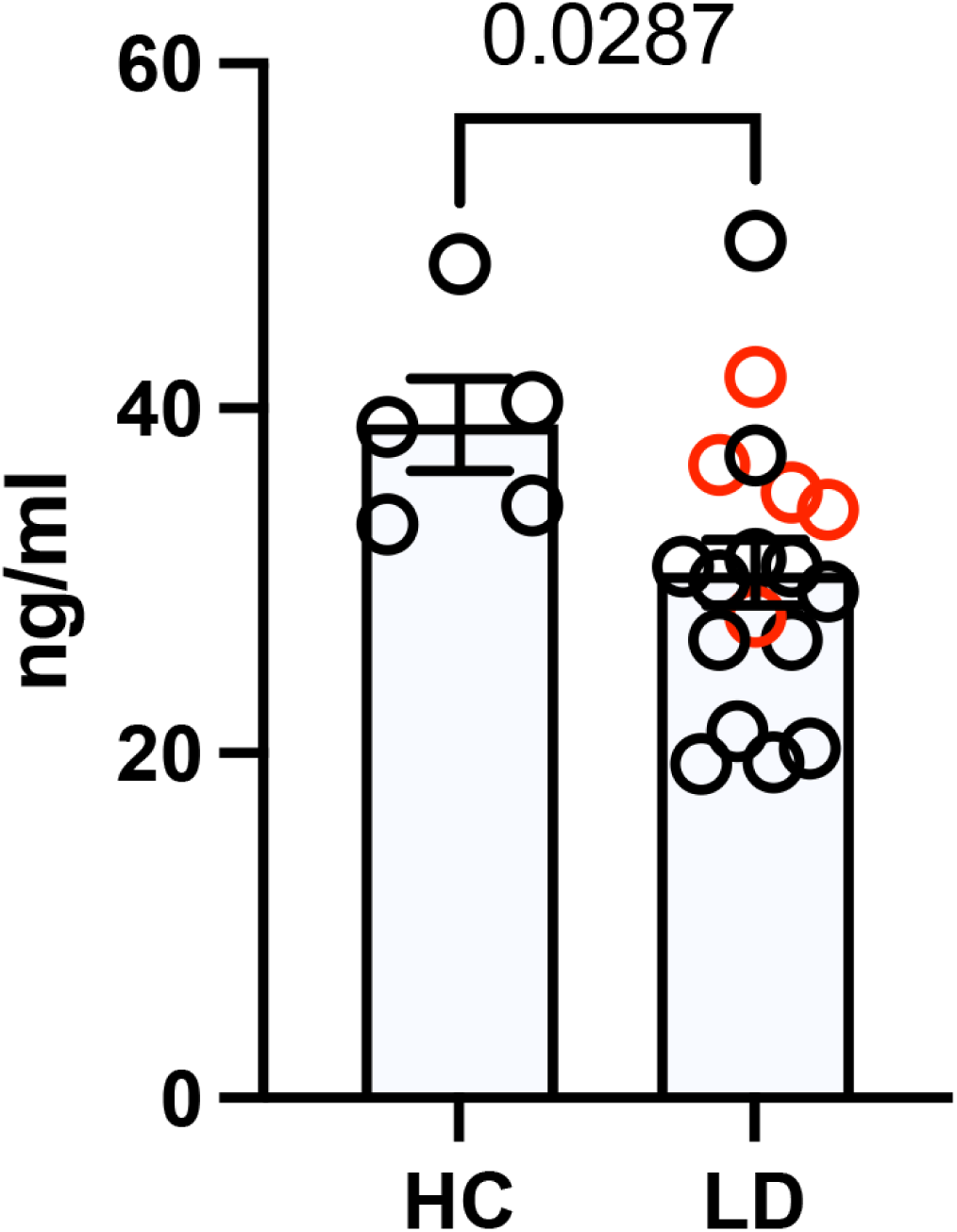
Serum SLPI levels in Lyme disease subjects versus healthy controls. The serum level of SLPI was measured by ELISA. Sera samples were from 5 adult healthy controls (HC). 18 samples were from people with Lyme disease (LD) including 5 samples from 3 subjects presenting with Lyme arthritis (red) and 13 samples from 4 subjects with erythema migrans (black). The error bar represents mean ± SEM and *p*-values are calculated using the non-parametric Mann-Whitney test.

### SLPI interacts with B. burgdorferi

It has been demonstrated that *B. burgdorferi* interacts with various mammalian proteins to establish infection in the mammalian host^40–43^. Thus, we postulated that *B. burgdorferi* could interact with SLPI to influence the progression of joint inflammation. To test this hypothesis, we probed a recently developed BASEHIT (**BA**cterial **S**election to **E**lucidate **H**ost-microbe **I**nteractions in high **T**hroughput) library with *B. burgdorferi*^40,44–46^. BASEHIT utilizes a genetically barcoded yeast display library expressing 3,324 human exoproteins, thus enabling a comprehensive screen of host-microbe interactions in a high-throughput fashion. Human SLPI is one of the exoproteins that passed the significance threshold, indicating *B. burgdorferi*-SLPI binding. To further establish that hSLPI directly binds to *B. burgdorferi*, we performed ELISA with whole cell *B. burgdorferi* lysates. We observed strong binding between whole cell *B. burgdorferi* lysates and hSLPI at a level as low as 10 nM (Figure 5A).

**Figure 5.**
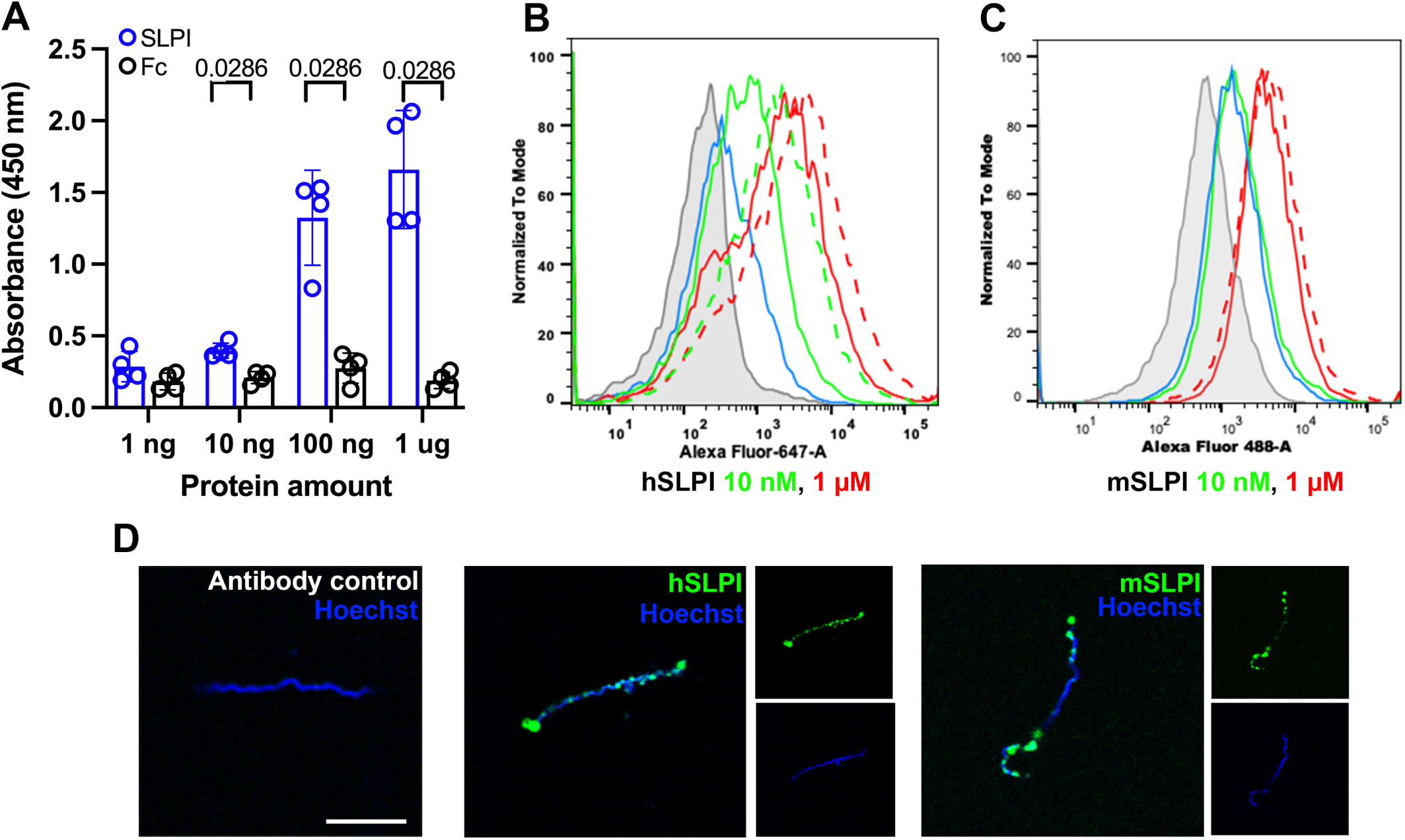
*B. burgdorferi* interaction with human and murine SLPI. (A) Sandwich ELISA results show the interaction of *B. burgdorferi* whole cell lysates with human SLPI. ELISA plates were coated with *B. burgdorferi* whole cell lysates and probed with increasing amount of human SLPI (blue) or human Fc proteins (black) as the negative control. The values plotted represent the mean ± SEM of duplicates from two experiments. *P*-value is displayed in the graph and determined using the non-parametric Mann-Whitney test. (B and C) Flow cytometry histograms show binding of human (B) and murine (C) SLPI to *B. burgdorferi* cultured at 33°C (solid line) and 37°C (dash line). *B. burgdorferi* was cultured to a density of 10^6^/ml. The same volume of cultures was incubated at 33°C or 37°C for 24 hrs before adding 10 nM (green) or 1 μM (red) human or murine SLPI. The binding was detected with goat anti human or murine SLPI and donkey anti goat AF647 or AF488. *B. burgdorferi* alone (grey) and antibody control (without SLPI, blue) were used as negative controls. (D) Immuno-fluorescent microscopy was used to directly observe the binding of *B. burgdorferi* with human and murine SLPI. Merged and single-color images are shown. Representative histograms and fluorescent images were shown from three independent experiments. Scale bar: 10 μm.

To extend these studies, flow cytometry was performed using intact *B. burgdorferi* and both human and murine SLPI. A significant increase in fluorescence intensity was observed when *B. burgdorferi*, cultivated at 33°C, was incubated with human SLPI at 10 nM and 1 µM level (Figure 5B). Though the binding to 10 nM rmSLPI was at background level, we observed a significant increase in fluorescence intensity when *B. burgdorferi* were incubated with 1 µM rmSLPI (Figure 5C). Flow cytometry also demonstrated increased binding of *B. burgdorferi* cultured at 37 °C to 1 µM hSLPI or rmSLPI (Figure 5, B and C). This indicates that the binding was more robust when performed at temperatures that *B. burgdorferi* encounter in the mammalian host. Immuno-fluorescent microscopy was an additional method that also demonstrated direct binding of *B. burgdorferi* with human or murine SLPI (Figure 5D).

In contrast to *B. burgdorferi* B31-A3, an infectious strain used throughout this study, we did not observe any binding between hSLPI and *B. burgdorferi* B31A, a high-passage non-infectious strain (Supplemental Figure 2A)^47,48^. The above observation further suggests that the direct interaction between SLPI and *B. burgdorferi* could impact the pathogenesis of murine Lyme arthritis. To further investigate the potential *B. burgdorferi* ligand that interacts with SLPI, we probed protease-treated *B. burgdorferi* lysates with hSLPI using ELISA. After treatment with proteinase K, we observed a marked decrease in binding of hSLPI to *B. burgdorferi* lysates (Supplemental Figure 2B). This result suggests that hSLPI can directly interact with a *B. burgdorferi* protein.

It has been showed that SLPI has antimicrobial effects against multiple gram-negative and positive bacteria^49–51^. However, using the BacTiter-Glow microbial cell viability assay, we did not observe any significant changes in *B. burgdorferi* viability in the presence of hSLPI (Supplemental Figure 3A). A previous study also demonstrated that the tick salivary protein, Salp15, specifically interacted with *B. burgdorferi* outer surface protein C (OspC)^52^. The binding of Salp15 protected spirochetes from killing by polyclonal mouse or rabbit antisera *in vitro*^52,53^. However, again, using the BacTiter-Glow microbial cell viability assay, the pre-incubation of hSLPI did not protect spirochetes from killing by mouse *B. burgdorferi* antisera (Supplemental Figure 3B). Thus, the importance of the SLPI-*B. burgdorferi* interaction and the direct effect of such an interaction on *B. burgdorferi* biology is likely independent of direct borreliacidal activity or any interference with the antibody-mediated *B. burgdorferi* killing.

## Discussion

Lyme arthritis has been extensively documented and studied in patients and *B. burgdorferi*-infected mice. The pathogenesis of Lyme arthritis is characterized by synovial tissue damage caused by infiltration of immune cells and excessive pro-inflammatory responses^7^. Transcriptomic studies also revealed the suppression of tissue repair genes in the synovial tissue of Lyme arthritis patients and tibiotarsal joint tissues of *B. burgdorferi*-infected mice^11,54^. However, the roles of the genes involved in tissue repair have not been studied.

SLPI strongly inhibits serine proteases, especially cathepsin G and elastase secreted by neutrophils^20^. The major function of SLPI is to prevent unnecessary tissue damage caused by excessive protease activity, thus promoting tissue repair and homeostasis^55^. The lack of SLPI impairs wound healing and tissue repair ^25^ and SLPI also inhibits NF-κB activation and downstream pro-inflammatory cytokine release from macrophages^27,29^. Thus, we hypothesized that SLPI plays an important role in Lyme arthritis. To test this hypothesis, we employed *SLPI*−/− C57BL/6 mice. As C57BL/6 mice only develop mild arthritis, if any, after challenge with *B. burgdorferi*, this mouse provided an ideal example to study whether the lack of SLPI could cause an arthritis-resistant mouse to become arthritis-susceptible. Compared to WT C57BL/6 mice, *B. burgdorferi* infection in the *SLPI*−/− mice consistently showed a significantly higher infection burden in tissues extracted from the ankle joint, which included periarticular structures (Figure 1E). Severe swelling and inflammation in the bursal and soft tissue around tibiotarsal joints were observed solely in the infected *SLPI*−/− mice (Figure 2). The significant increase in the infection burden in *SLPI*−/− mice can contribute to the enhanced periarticular inflammation following *B. burgdorferi* infection. However, these data also suggest the importance of SLPI in controlling the development of inflammation in periarticular tissues of *B. burgdorferi*-infected mice. Indeed, in a *Streptococcal* cell wall (SCW)-induced arthritis model in rats, the intraperitoneal injection of SLPI significantly decreased the severity of joint swelling^34^. Targeting the SLPI-associated anti-protease pathways could also potentially be a strategy for ameliorating periarticular inflammation that occurs in some rheumatic diseases.

Analysis of *B. burgdorferi* infection in the *SLPI*−/− and WT mice revealed a significant increase in infiltrating neutrophils and macrophages in the periarticular tissues of *SLPI*−/− mice (Figure 3, A and B). This observation is consistent with clinical studies that showed a high percentage of neutrophils in the synovial fluid during active infection^56,57^. In post-treatment Lyme arthritis, fewer neutrophils and more macrophages are present in patients’ synovial fluid^7^. In arthritis-susceptible C3H/He mice, *B. burgdorferi* infection also leads to neutrophil infiltration in the periarticular tissues as well as in the synovium of ankle joints^13^. The neutrophil chemoattractant KC (CXCL1) and receptor CXCR2 mediates neutrophil recruitment and is critical for the development of murine Lyme arthritis. Both *KC*−/− and *CXCR2*−/− C3H mice developed significantly less ankle swelling when infected with *B. burgdorferi*^16,18^. Consistently, we observed a significant increase in CXCR2 gene expression in the tibiotarsal joint tissues (Figure 3C), which can recruit neutrophils and cause more severe inflammatory soft tissue infiltrates in the *SLPI*−/− mice. Monocyte chemoattractant protein-1 (MCP-1/CCL2) and receptor CCR2 contribute to macrophage infiltration^58^. Though little to no difference in arthritis was observed in the CCR2−/− mice, a high level of MCP-1 was detected in the tibiotarsal tissues of *B. burgdorferi*-infected, arthritis-susceptible C3H/He mice, suggesting a function for macrophages^18^. In the infected *SLPI*−/− mice, a significant increase in both MCP-1 and CCR2 gene expression was observed in the tibiotarsal tissues (Figure 3, D and E). Our data suggest that both neutrophils and macrophages contribute to the severe periarticular inflammation in the infected *B. burgdorferi-*infected *SLPI*−/− mice.

To investigate the underlying mechanism whereby SLPI deficiency enhanced periarticular joint inflammation, we examined the serum cytokine/chemokine profile of *B. burgdorferi*-infected *SLPI*−/− and WT mice. There was a significant increase in IL-6 solely in the infected *SLPI*−/− mice (Figure 3F). An elevated IL-6 level has been demonstrated in the serum, synovial fluid, and synovial tissue from Lyme arthritis patients^9,56^. IL-6 is also pivotal in the pathogenesis of rheumatoid arthritis and correlates with the disease severity and joint destruction^37^. It has been shown that IL-6 recruited neutrophils in an *in vitro* co-culture rheumatoid arthritis model^59^. Neutrophils can be activated by IL-6 through binding of IL-6 receptor (IL-6R)^37^. Activated neutrophils release several neutrophil serine proteases (NSPs) including neutrophil elastase (NE), cathepsin G (CTSG), and proteinase-3 (PR3), which can lead to potent cartilage destruction^31^. Indeed, we observed a significant increase in the NE level in the serum of *B. burgdorferi*-infected *SLPI*−/− but not WT mice (Figure 3G). In the arthritis susceptible C3H/HeN mice, *B. burgdorferi* infection also induced a significant increase in the serum NE level (Figure 3H). Interestingly, despite the known function of TNF-α in inflammatory responses, we did not observe any significant changes in either TNF-α serum protein levels or TNF-α gene expression levels in the *B. burgdorferi*-infected *SLPI*−/− and WT mice (Supplemental Figure 4). The result is consistent with previous microarray data that did not show significant changes in TNF-α levels in the C57BL/6 mice following *B. burgdorferi* infection^54^. The above data indicate that the excessive serum neutrophil elastase contributed to the periarticular inflammation in the *B. burgdorferi*-infected *SLPI*−/− mice.

Matrix metalloproteinases (MMPs) target extracellular matrix and cause articular cartilage destruction^60^. A correlation between IL-6 and MMPs expression has been reported in the context of rheumatoid arthritis^38,61^. Elevated levels of several MMPs have also been found in the synovial fluid of Lyme arthritis patients^39^. Thus, we also investigated the MMPs profile in the *B. burgdorferi*-infected *SLPI*−/− and WT mice. We observed a significant increase in the serum level of MMP-8 in the infected *SLPI*−/− mice (Figure 3I). MMP-8 is known as neutrophil collagenase^62^. Using synovial fluid samples, it has been reported that the level of MMP-8 was significantly higher in the patients with post infectious Lyme arthritis than patients with active infection^63^. A comprehensive examination of the MMP profile in the synovial fluid of patients with Lyme arthritis revealed elevated levels of MMP-1, −3, −13, and −19^64^. *B. burgdorferi* infection induced MMP-3 and MMP-19 in the C3H/HeN mice but not in the Lyme arthritis-resistant C57BL/6 mice^64^. The differences in the MMP profiles provide an explanation for the differences between human and murine Lyme arthritis. This finding further emphasizes that excessive protease activity can contribute to the severity of periarticular inflammation in *B. burgdorferi*-infected mice.

Previous research using serum, synovial fluid and tissue from Lyme arthritis patients has been heavily focusing on innate and adaptive immune responses^7^. As a result, limited data can be found regarding anti-protease and protease responses during Lyme arthritis in human patients. In this study, we tested serum SLPI level in 5 healthy subjects, 8 Lyme disease patients, 3 of whom had overt arthritis. Though the number of healthy subjects is small, the median level of SLPI tested here (38.92 ng/ml, Figure 4) is comparable with previous studies showing in average about 40 ng/ml SLPI in the serum from healthy volunteers ^65,66^. While the clinical manifestation of 5 of the patients with Lyme disease was an EM skin lesion (Supplemental Table 1), some symptom persisted several months after diagnosis, a timeframe when acute arthritis often develops. We observed decreased SLPI in the serum of these patients (Figure 4), suggesting an inverse relationship between the SLPI level and *B. burgdorferi*-infection. However, a large number of sera and synovial fluid samples from patients with Lyme arthritis and other clinical manifestations of Lyme disease are needed to establish a definitive association.

*B. burgdorferi* first infects the skin of a vertebrate host following a tick bite, then disseminates throughout the body, colonizes various tissue, evades immune responses, and persists for a significant period of time. To survive the above processes, *B. burgdorferi* interacts with various mammalian proteins, including decorin^41^, fibronectin^42^, and plasminogen^43^, among others. To comprehensively study the potential interaction between *B. burgdorferi* and the host, our lab employed the BASEHIT (**BA**cterial **S**election to **E**lucidate **H**ost-microbe **I**nteractions in high **T**hroughput) to assess the interactions between *B. burgdorferi* and 3,336 human extracellular and secreted proteins^44,46^. Using BASEHIT, our lab previously identified a strong interaction between *B. burgdorferi* and Peptidoglycan Recognition Protein 1 (PGYRP1)^40^. Increased infection burden in the heart and joint was observed in the mice lacking PGYRP1, suggesting a role of PGYRP1 in the host response to *B. burgdorferi* infection. Expanding the use of BASEHIT, CD55 was identified to bind *Borrelia crocidurae* and *Borrelia persica*, two pathogens causing relapsing fever^45^. CD55-deficient mice infected with *B. crocidurae* displayed lower pathogen load and elevated pro-inflammatory cytokines. The above data demonstrate BASEHIT as an effective method to identify host factors important in *B. burgdorferi* pathogenesis *in vivo*. In this study, we identified an interaction between SLPI and *B. burgdorferi* using BASEHIT library screening and subsequent flow cytometric analysis (Figure 5). The antimicrobial activity of SLPI has been demonstrated against both gram-negative and positive bacteria including *Escherichia coli*, *Pseudomonas aeruginosa*^49^, *Mycobacteria tuberculosis*^50^, *Staphylococcus aureus*^51^, and *Staphylococcus epidermidis*^49^. Interaction between the positive charges of SLPI and the negative charges of bacteria surface, including lipopolysaccharide (LPS), can destabilize bacteria cell wall leading to the bactericidal effect^67^. *B. burgdorferi* do not have LPS^68^ and this may account for the absence of the bactericidal effect of SLPI against *B. burgdorferi* (Supplemental Figure 3A). Future research is needed to understand the significance of the SLPI-*B. burgdorferi* binding in the development of periarticular inflammation. The potential *B. burgdorferi* protein that interact with SLPI remains unknown. It is our hypothesis that SLPI may bind and inhibit an unknown *B. burgdorferi* virulence factor that could contribute to the development of murine Lyme arthritis.

In conclusion, our data demonstrated the importance of SLPI in suppressing *B. burgdorferi*-induced periarticular inflammation in mice by inhibiting recruitment of neutrophils and macrophages and subsequent protease levels. We propose that, during the active infection of the murine joint structures, the binding of *B. burgdorferi* with SLPI depletes the local environment of SLPI. Such binding is specific to the infectious strain of *B. burgdorferi.* As a potent anti-protease, the decrease in SLPI results in excessive protease activity, including neutrophil elastase and MMP-8. These unchecked proteases can lead to extensive tissue inflammation. Our study is the first to emphasize the importance of an anti-protease-protease balance in the development of the periarticular inflammation seen in *B. burgdorferi*-infected mice.

## Materials and methods

### Sex as a biological variable

Females *SLPI*−/− and WT C57BL/6 mice were used for the *in vivo B. burgdorferi* infection. We have examined *B. burgdorferi* infection in both male and female C57BL/6 mice and no differences in the development of infection or disease have been noted. Both male and female Lyme disease patients were included in the study. Sex was not considered as a biological variable.

### Study approval

This study used archived serum samples from adult Lyme disease subjects and healthy controls that were previously collected under NIH U19AI089992 with approval of the Yale University Institutional Review Board for human subjects research (IRB protocol# 1112009475). All the animal experiments in this study were performed in accordance with the Guidelines for the Care and Use of Laboratory Animals of the National Institutes of Health. The animal protocols were approved by the Institutional Animal Care and Use Committee at the Yale University School of Medicine.

### Measurement of serum SLPI levels in Lyme disease subjects and controls

SLPI levels were measured in a total of 23 serum samples from 7 subjects at the time of Lyme disease diagnosis (4 with a single erythema migrans lesion and 3 with the late manifestation of Lyme arthritis) and from 5 healthy controls. Serum samples from Lyme disease subjects were available at up to 3 times points: 1) study entry, range 0-9 days after onset of symptoms, 2) 30 days post diagnosis, and 3) up to 3 months after completion of antibiotic therapy (range 4.5 – 6 months after dx). Additional details can be found in Supplemental Table 1. The level of SLPI in the serum was measured using the Human SLPI DuoSet ELISA kit (R&D Systems, #DY1274-05).

### B. burgdorferi culture

*B. burgdorferi* B31-A3, an infectious clonal derivative of the sequenced strain B31, was a generous gift from Dr. Utpal Pal at the Department of Veterinary Medicine, University of Maryland, College Park^69^. *B. burgdorferi* B31-A3 and *B. burgdorferi* B31A was grown in Barbour-Stoenner-Kelly H (BSK-H) complete medium (Sigma-Aldrich, #B8291) in a 33°C setting incubator. The live cell density was determined by dark field microscopy and using a hemocytometer (INCYTO, #DHC-N01). Low passage (P<3) *B. burgdorferi* B31-A3 was used throughout this study.

### *In vivo* infection of mice

The *SLPI*−/− C57BL/6 mice have been described previously^70,71^. The wild-type (WT) C57BL/6 mice were purchased from the Jackson Laboratory and used as the controls. 5 to 7 weeks of age female WT and *SLPI*−/− C57BL/6 mice were used for infection. 4 to 6 weeks of age female C3H/HeN mice were purchased from Charles River Laboratories and used for infection. Both C57BL/6 and C3H/HeN mice were infected with low passage 10^5^ *B. burgdorferi* subcutaneously (5–9 mice/group). PBS sham-infected mice were used as controls. Mice were euthanized approximately 3-week post infection within a 3-day window (between 21 to 24 dpi) based on the feasibility and logistics of the laboratory. Ear punch biopsies were taken at 7-, 14- and between 21 to 24-day post-infection (dpi) to determine the infection burden in the skin. Between 21 to 24 dpi, mice were euthanized, and heart and joint tissues were collected to quantify the spirochete burden. The protocol for the use of mice was reviewed and approved by the Yale Animal Care and Use Committee.

### Quantification of *Borrelia* burden

DNA was extracted from heart, tibiotarsal joint, and ear punch samples using Qiagen DNeasy Blood & Tissue Kit, Qiagen. Quantitative PCR was performed using iQ-SYBR Green Supermix (Bio-Rad). For quantitative detection of *B. burgdorferi* burden within mouse tissue samples, q-PCR was performed with DNA using flagellin (*flaB*), a marker gene for *Borrelia* detection. The mouse β*-actin* gene was used to normalize the amount of DNA in each sample. The nucleotides sequences of primers used in specific PCR applications are described previously^40^.

### Joint histopathology analysis

Mice were euthanized by CO_2_ asphyxiation and one rear leg from each mouse were dissected, immersion-fixed in Bouin’s solution (Sigma-Aldrich, #HT10132). Fixed tissues were embedded, sectioned, and stained with hematoxylin and eosin (HE) by routine methods (Comparative Pathology Research Core in the Department of Comparative Medicine, Yale School of Medicine). Periarticular and joint inflammation was scored in a blinded fashion in a graded manner from 0 (negative), 1 (minimal), 2 (moderate), to 3 (severe).

### Flow cytometry to quantify infiltrating cells in joint tissues in mice

The WT and *SLPI*−/− C57BL/6 mice were infected with *B. burgdorferi* as described above. The mice were euthanized between 21 to 24 dpi. The ankle joints were cut out at around 0.7 cm proximal to the ankle joint. The portion distal to the midfoot was discarded, and the skin removed. The bone marrow cells were flushed out with RPMI 1640 (Gibco, #11875-093) using a 27-gauge needle. The bone marrow-depleted ankles were cut into 3-4 mm sized tissue pieces and incubated with digestion media containing 2.4 mg/ml hyaluronidase (Sigma-Aldrich, #H3506), 1 mg/ml collagenase (Sigma-Aldrich, #C2139) in RPMI 1640 (Gibco, #11875-093) supplemented with 10% fetal bovine serum (FBS) for 1 h at 37 °C with 5%CO_2_. The digestion media containing the tissue pieces were passed through a 70 μm cell strainer (Thermo Scientific, #352350). The remaining tissue pieces were mashed using a 10 ml syringe plunger. The digestion media containing the isolated cells were neutralized with RPMI 1640 with 10% FBS^35^. The red blood cells were lysed by ACK Lysing buffer (Gibco, #A1049201). The cells were rinsed and resuspended in FACS buffer and ready for staining for flow cytometry.

The cells were incubated with Fc receptor antibody (TruStain FcXTM anti-mouse CD16/32) (BioLegend, #101320), and antibodies including PerCP anti-mouse CD45 (BioLegend, #103130), BV711 anti-mouse Ly6G (BioLegend, #127643), PE anti-mouse CD11b (BioLegend, #101208), APC/CY7 anti-mouse CX3CR21 (BioLegend, #149047), FITC anti-mouse Ly6C (BioLegend, #128005), APC anti-mouse CD64 (BioLegend, #139305) and LIVE/DEADTM fixable violet stain kit (Invitrogen, #L34955) on ice for 30 min. The samples were rinsed twice with FACS buffer and run through BD LSRII (BD bioscience). The data was then analyzed by FlowJo^36^.

### Gene expression evaluation by quantitative real-time PCR

Mice were euthanized between 21 to 24 dpi. The ankle joints were excised as described above, snap-frozen in liquid nitrogen, and stored at −80°C. The frozen tissue was pulverized in liquid nitrogen using a mortar and pestle^72^. The RNA was purified using Trizol (Invitrogen, #15596-018) following a published protocol^73^. cDNA was synthesized using the iScript cDNA Synthesis Kits (Bio-Rad, #1708891). qPCR was performed using iQ SYBR Green Supermix (Bio-Rad, #1725124). The relative expression of each target gene was normalized to the mouse β*-actin* gene. The target genes and corresponding primer sequences are shown in the Key Resource Table.

### Murine neutrophil elastase, cytokine, chemokine, and MMP profile

Blood samples from each group of mice was collected by cardiac puncture between 21 to 24 dpi and sera were collected. The murine neutrophil elastase level was measured using the Mouse Neutrophil Elastase/ELA2 DuoSet ELISA (RnD Systems, #DY4517-05). Serum was sent for cytokine analysis by the Mouse Cytokine/Chemokine 32-Plex Discovery Assay Array (MD32), and the Mouse MMP 5-Plex Discovery Assay Array (MDMMP-S, P) performed by Eve Technologies. The cytokines and chemokines represented by MD32 are Eotaxin, G-CSF, GM-CSF, IFN-γ, IL-1α, IL-1β, IL-2, IL-3, IL-4, IL-5,IL-6, IL-7, IL-9, IL-10, IL-12 (p40), IL-12 (p70), IL-13, IL-15, IL-17A, IP-10, KC, LIF, LIX, MCP-1, M-CSF, MIG, MIP-1α, MIP-1β, MIP-2, RANTES, TNFα, and VEGF. The MMPs represented by MDMMP-S, P are MMP-2, MMP-3, MMP-8, proMMP-9, and MMP-12.

### Purification of recombinant murine SLPI

The murine *SLPI* cDNA ORF clone was purchased from GenScript (OMu22721). The coding sequence was subsequently cloned into pET22b(+) expression vector (Novagen) in frame with the pelB signal peptide using Gibson Assembly^74^. *E. coli* strain Rosetta-gami 2 (DE3) (Novagen, #71351) was transformed with the SLPI-pET22b+ and grown at 37°C with ampicillin (100 μg/ml), tetracycline (12.5 μg/ml), streptomycin (50 μg/ml) and chloramphenicol (34 μg/ml). Cells were induced with 1 mM IPTG (18°C, overnight), harvested, and lysed with BugBuster Protein Extraction Reagent (Novagen, #70921-3). Recombinant mSLPI was purified with a Ni-NTA resin column as described by the manufacturer (Qiagen). To evaluate the activity of the purified rmSLPI, the trypsin inhibitory activity was assayed with the fluorescent substrate Mca-RPKPVE-Nval-WRK(Dnp)-NH2 Fluorogenic MMP Substrate (R&D Systems, #ES002) and the absorbance was monitored at 405 nm using a fluorescent plate reader (Tecan).

### Flow cytometry to validate *B. burgdorferi*-SLPI binding

Actively growing low passage *B. burgdorferi* was cultured to a density of 10^6^–10^7^ cells/ml and harvested at 10,000 g for 10 min. Cells were rinsed twice with PBS and blocked in 1%BSA for 1 h at 4°C. The cells were pelleted, rinsed, resuspended, and incubated with 10 nM and 1 µM human SLPI (R&D Systems, #1274-PI-100) and murine SLPI (produced in lab as described above) at 4°C for 2 h. The binding was detected with goat anti human or murine SLPI (R&D Systems, #AF1274 and AF1735) and Alexa Fluor 488 or Alexa Fluor 647 donkey anti goat IgG (H+L) (Invitrogen, #A32814, A-21447). The samples were fixed with 2% PFA before running through BD LSRII Green (BD bioscience). The data was then analyzed by FlowJo. The integrity of *B. burgdorferi* organism was confirmed by Hoechst 33324/ propidium iodide double staining (Supplemental Figure 5). The fixed *B. burgdorferi* sample was included as a positive control for the propidium iodide staining. Permeabilization was not performed during this protocol. Thus, the binding detected was on the bacterial outer surface.

### ELISA to validate *B. burgdorferi*-SLPI binding

*B. burgdorferi* was cultured to a density of 10^6^–10^7^ cells/ml and harvested at 10,000 g for 10 min. To make the *B. burgdorferi* lysate, cells were rinsed twice with PBS, pelleted, and lysed using BugBuster Protein Extraction Reagent (Novagen, #70921-3). Protein concentration in the lysate was measured by absorbance at 280 nm using the nanodrop (Fisher Scientific). For the protease assay, the *B. burgdorferi* lysate was incubated in the presence or absence of proteinase K (0.2 mg/ml, Thermo Scientific, #EO0491) for 10 min. In an immuno 96-well plate (MaxiSorp), wells were coated with 200 ng of *B. burgdorferi* lysate. Samples were blocked with 1% BSA followed by incubation with human SLPI at varying concentrations (1–1000 ng) for 1 h at room temperature. The binding was probed with goat anti human SLPI (R&D Systems, #AF1274) and rabbit anti goat IgG (whole molecule)-Peroxidases antibody (Sigma-Aldrich, #A8919-2ML). KPL Sureblue TMB Microwell Peroxidase substrate, 1-component (Seracare, #5120-0077) was used. The reaction was stopped with KPL TMB stop solution (Seracare, #5150-0021), and absorbance was read at 450 nm.

### Immunofluorescence assay

Actively growing low passage *B. burgdorferi* was cultured to a density of 10^6^–10^7^ cells/ml, rinsed twice with PBS and blocked with 1% BSA for 1 h at 4°C. *B. burgdorferi* was incubated with human or murine SLPI at 4°C for 2 h. The spirochetes were probed with goat anti human or murine SLPI (R&D Systems, #AF1274 and AF1735) and Alexa Fluor 488 donkey anti goat IgG (H+L) (Invitrogen, #A32814). *B. burgdorferi* were then stained with Hoechst 33342 (Invitrogen, #H1399). The samples were fixed with in 2% PFA before imaged with Leica SP8. The integrity of *B. burgdorferi* organism was confirmed in Supplemental Figure 5. Permeabilization was not performed during this protocol. Thus, the binding detected was on the bacterial outer surface.

### The BacTiter Glo microbial cell viability assay to quantify *B. burgdorferi* viability

The BacTiter Glo microbial cell viability assay quantifies the ATP present in the microbial culture by measuring luminescence. The amount of ATP is proportional to the number of viable cells in culture^40,45,75^. To test the borreliacidal activity of human SLPI, 1×10^5^ spirochetes were treated with 0–10 μM hSLPI (R&D Systems, #1274-PI-100) at 33°C for 48 h. The luminescence was measured using a fluorescence plate reader (Tecan). The percent viability was normalized to the control spirochetes culture without hSLPI treatment. To test the effect of hSLPI on the antibody-mediated *B. burgdorferi* killing, 1×10^5^ spirochetes were pretreated with 0–5 μM hSLPI (R&D Systems, #1274-PI-100) at 33°C for 2 h. 20% mouse *B. burgdorferi* antisera were then added and incubated for 2 and 4 h. The mouse antisera were collected from *B. burgdorferi* infected mice between 21 to 24 dpi. The luminescence was measured as described above. The percent viability was normalized to the control spirochetes culture without any treatment.

### Statistical analysis

The analysis of all data was performed using the non-parametric Mann-Whitney, or ANOVA using Prism 10 software (GraphPad Software, Inc., San Diego, CA). A *p*-value of <0.05 was considered statistically significant.

## ACKNOWLEGMENTS

We are grateful to Dr. Narasimhan for her input and suggestions during experiment design, and to Dr. Ming-Jie Wu for his assistance in conducting experiments. We are grateful to Ms. Ming Li for her effort in preparing human sera samples. This work was supported by NIH grants AI165499 and AI138949, the Steven and Alexandra Cohen Foundation, and the Howard Hughes Medical Institute Emerging Pathogens Initiative.

## AUTHOR CONTRIBUTIONS

QY and EF designed, performed, and analyzed experiments and draft the manuscript. XT designed and performed experiments. TH performed the BASEHIT library screen. RH scored the severity of murine ankle inflammation. AB and LB provided sera from Lyme disease patients and edited the manuscript. AN provided the *SLPI*−/− mice. AR developed the BASEHIT screen. EF scored the visual swelling level of murine tibiotarsal joints and supervised the research.

## Figure Legends

**Supplemental Figure 1. The macrophages population analyzed using Ly6G-negative gating strategy.** The macrophage population was first gated on the Ly6G-negative population, then gated on the CD64 positive cells among the CX3CR1 positive myeloid cells. Results from two independent experiments were pooled and shown here.

**Supplemental Figure 2. The binding of human SLPI to non-infectious *B. burgdorferi B31A* and proteinase K-treated *B. burgdorferi B31A3*.** (A) Flow cytometry histogram shows the lack of binding of human SLPI (1 μM, red) to non-infectious *B. burgdorferi B31A. B. burgdorferi* alone (grey) and antibody control (without SLPI, blue) were used as negative controls. A representative histogram from two independent experiments was shown. (B) ELISA result shows the interaction between human SLPI and *B. burgdorferi B31A3* whole cell lysates in the presence (blue) or absence (black) of proteinase K. ELISA plates were coated with *B. burgdorferi B31A3* whole cell lysates and probed with increasing amount of human SLPI. The values plotted represent the mean ± SEM of triplicates from one experiment.

**Supplemental Figure 3. The effect of human SLPI binding on *B. burgdorferi* viability and antibody-mediated killing.** (A) Human SLPI (hSLPI, 0-10 μM) was incubated with 10^5^ *B. burgdorferi* at 33°C for 48 hours. The viability was assessed by BacTiter Glo microbial cell viability assay. The percent viability was normalized to the control spirochetes culture without hSLPI treatment. Results from one independent experiment performed in triplicate samples are shown here. (B) Human SLPI (hSLPI, 0-5 μM) was incubated with 10^5^ *B. burgdorferi* at 33°C for 2 hours. 20% mouse *B. burgdorferi* antisera were then added for 2 and 4 h. The viability was measured as described above. The percent viability was normalized to the control spirochetes culture without any treatment. Results from two independent experiments performed in duplicate samples are shown here.

**Supplemental Figure 4. Serum and gene expression levels of TNF-α.** (A) The serum level of TNF-α was assessed using a mouse cytokine/chemokine 32-plex array. (B) The gene expression level of TNF-α was assessed in the tibiotarsal tissue using RT-qPCR. Serum was obtained by cardiac puncture and the tibiotarsal tissue was collected from WT and *SLPI*−/− C57BL/6 mice with/without *B. burgdorferi* infection between 21 to 24 dpi. black, PBS-sham infection; red, *B. burgdorferi* infection. Each data point represents an individual animal. The error bar represents mean ± SEM and *p*-values, and were calculated using the non-parametric Mann-Whitney test.

**Supplemental Figure 5. Hoechst 33324 and propidium iodide double staining of *B. burgdorferi* whole organism with and without fixation.** Immuno-fluorescent microscopy was used to directly observe the staining of *B. burgdorferi*. Merged and single-color images are shown. Representative images were shown from two independent experiments. Scale bar: 10 μm.

**Supplemental Table 1. Subject Characterization**

**Supplemental Table 2. The primers used in this study.**

## Key resources table

| Reagent type (species) or resource | Designation | Source or reference | IDENTIFIER |
| --- | --- | --- | --- |
| Antibodies | TruStain FcXTM anti-mouse CD16/32 | BioLegend | Cat#101320 |
| Antibodies | PerCP anti-mouse CD45 | BioLegend | Cat#103130 |
| Antibodies | BV711 anti-mouse Ly6G | BioLegend | Cat#127643 |
| Antibodies | PE anti-mouse CD11b | BioLegend | Cat#101208 |
| Antibodies | APC/CY7 anti-mouse CX3CR21 | BioLegend | Cat#149047 |
| Antibodies | FITC anti-mouse Ly6C | BioLegend | Cat#128005 |
| Antibodies | APC anti-mouse CD64 | BioLegend | Cat#139305 |
| Antibodies | goat anti human SLPI | R&D Systems | Cat#AF1274 |
| Antibodies | goat anti murine SLPI | R&D Systems | Cat#AF1735 |
| Antibodies | Alexa Fluor 488 donkey anti goat IgG (H+L) | Invitrogen | Cat#A32814 |
| Antibodies | Alexa Fluor 647 donkey anti goat IgG (H+L) | Invitrogen | Cat#A-21447 |
| Bacterial strains | *Borrelia. burgdorferi* B31-A3 | Dr. Utpal Pal's laboratory | N/A |
| Bacterial strains | *Borrelia. burgdorferi* B31-A | This manuscript | N/A |
| Bacterial strains | *E. coli* Rosetta-gami 2 (DE3) competent cell | Novagen | Cat#71351 |
| Biological samples | Mouse tibiotarsal tissue | This manuscript | N/A |
| Recombinant proteins | human SLPI | R&D Systems | Cat#1274-PI-100 |
| Chemical | iQ SYBR Green Supermix | Bio-Rad | Cat#1725124 |
| Chemical | Barbour-Stoenner-Kelly H (BSK-H) complete medium | Sigma-Aldrich | Cat#B8291 |
| Chemical | Bouin's solution | Sigma-Aldrich | Cat#HT10132 |
| Chemical | Hyaluronidase | Sigma-Aldrich | Cat#H3506 |
| Chemical | Collagenase | Sigma-Aldrich | Cat#C2139 |
| Chemical | ACK Lysing buffer | Gibco | Cat#A1049201 |
| Chemical | Trizol | Invitrogen, | Cat#15596-018 |
| peptide | Mca-RPKPVE-Nval-WRK(Dnp)-NH2<br>Fluorogenic MMP Substrate | R&D Systems | Cat#ES002 |
| Chemical | BugBuster Protein Extraction Reagent | Novagen | Cat#70921-3 |
| Chemical | Proteinase K | Thermo Scientific | Cat#EO0491 |
| Chemical | Ni-NTA agarose | QIAGEN | Cat#30230 |
| Chemical | KPL Sureblue TMB Microwell Peroxidase substrate, 1-component | Seracare | Cat#5120-0077 |
| Chemical | KPL TMB stop solution | Seracare | Cat#5150-0021 |
| Chemical | Hoechst 33342 | Invitrogen | Cat#H1399 |
| Chemical | RPMI 1640 | Gibco | Cat#11875-093 |
| Critical commercial assays | LIVE/DEAD<sup>TM</sup> fixable violet stain kit | Invitrogen | Cat#L34955 |
| Critical commercial assays | iScript cDNA Synthesis Kit | Bio-Rad | Cat#1708891 |
| Critical commercial assays | DNeasy Blood & Tissue Kit | Qiagen |  |
| Critical commercial assays | Gibson Assembly Kit | NEB |  |
| Critical commercial assays | Mouse Neutrophil Elastase/ELA2 DuoSet ELISA | RnD Systems | Cat#DY4517-05 |
| Critical commercial assays | Human SLPI DuoSet ELISA | RnD Systems | Cat#DY1274-05 |
| Critical commercial assays | BacTiter-Glo Microbial Cell Viability Assay kit | Promega | Cat#G8230 |
| Critical commercial assays | Mouse MMP 5-Plex Discovery Assay Array (MDMMP-S, P) | Eve Technologies | N/A |
| Critical commercial assays | Mouse Cytokine/Chemokine 32-Plex Discovery Assay Array (MD32) | Eve Technologies | N/A |
| Experimental models: Organisms/strains | *SLPI*<sup>-/-</sup> mice | Dr. Akira Nakamura's laboratory<sup>68, 69</sup> | https://doi.org/10.3389/fimmu.2017.01538<br>https://doi.org/10.1084/jem.20021824 |
| Experimental models: Organisms/strains | C3H/HeN mice | Charles River Laboratories | N/A |
| Experimental models: Organisms/strains | C57BL/6 mice | Jackson Laboratory | Stock #: 000664; RRID: IMSR_JAX:000664 |
| Recombinant DNA | Plasmid: Murine *SLPI* cDNA ORF clone | GenScript | OMu22721 |
| Recombinant DNA | Plasmid: pET-22b (+) | Novagen | Cat#69744-3 |
| Software and algorithms | Prism | GraphPad | RRID:SCR_002798 |
| Software and algorithms | FlowJo | BD Biosciences | https://www.flowjo.com/ |

