## Supplementary figures and images for "Secretory leukocyte protease inhibitor influences periarticular joint inflammation in *B. burgdorferi*-infected mice"

### Supplemental Figures

#### Supplemetal Figure 1

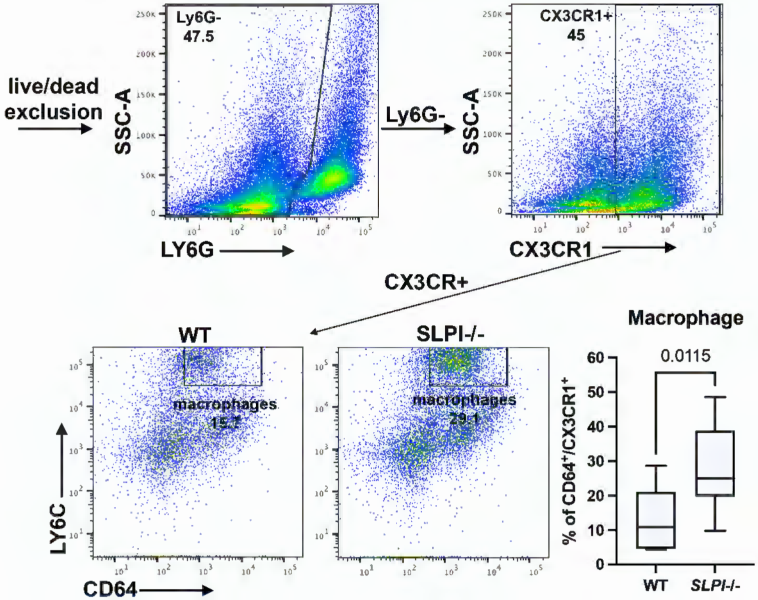

#### Supplemetal Figure 2

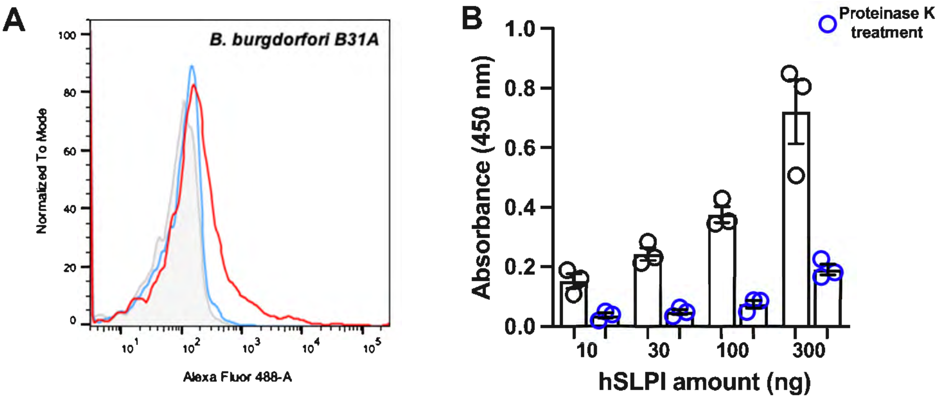

#### Supplemetal Figure 3

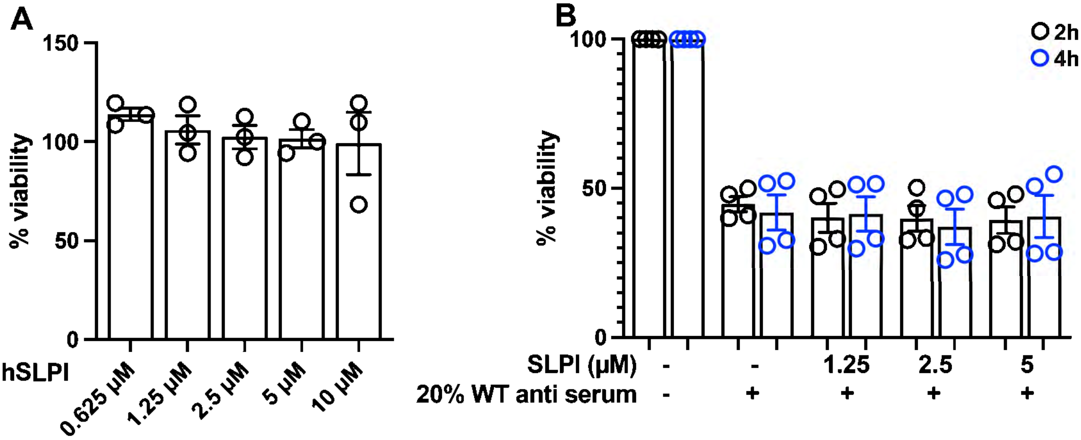

#### Supplemetal Figure 4

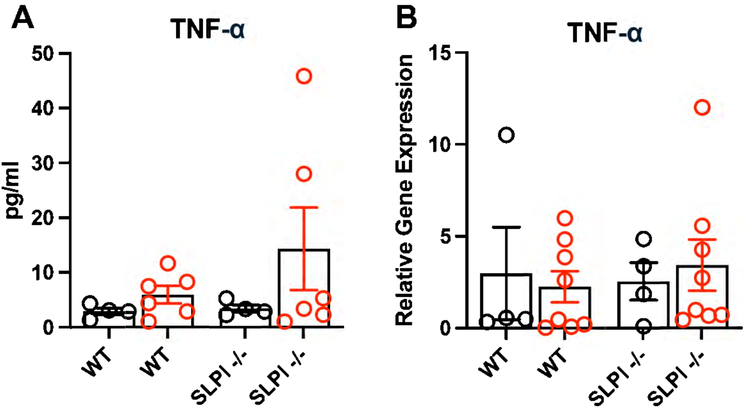

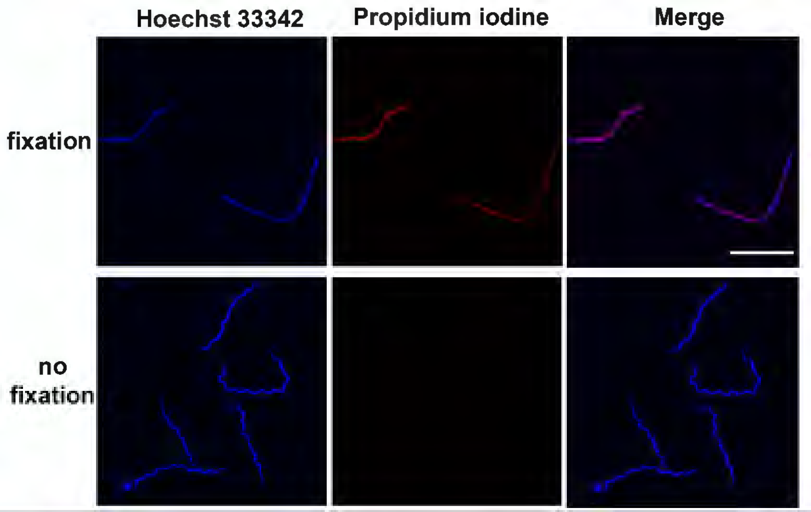
