## Supplementary material for "Secretory leukocyte protease inhibitor influences periarticular joint inflammation in *B. burgdorferi*-infected mice": Supplmental Table 1

**Supplemental Table 1.**

**Subject Characterization**

| Disease stage | Age | Gender | Disease Presentation | Time point of Sera Collection |
| --- | --- | --- | --- | --- |
| Acute Localized  Infection | 68 | F | Single EM | 0 |
|  |  |  |  | 1 |
|  |  |  |  | 2 |
| Acute Localized  Infection | 63 | F | Single EM | 0 |
|  |  |  |  | 1 |
|  |  |  |  | 2 |
| Acute Localized  Infection | 65 | F | Single EM | 0 |
|  |  |  |  | 1 |
|  |  |  |  | 2 |
| Acute  Localized  Infection | 63 | F | Single EM | 0 |
|  |  |  |  | 1 |
|  |  |  |  | 2 |
| Late Disease | 45 | M | Arthritis | 0 |
|  |  |  |  | 1 |
| Late Disease | 35 | M | Arthritis | 1 |
| Late Disease | 67 | M | Arthritis | 0 |
|  |  |  |  | 1 |
| Healthy Control | 74 | F | NA | NA |
| Healthy Control | 65 | F | NA | NA |
| Healthy Control | 66 | M | NA | NA |
| Healthy Control | 73 | M | NA | NA |
| Healthy Control | 59 | M | NA | NA |

Demographic information at the time of initial sample collection. Time points: 0=time of diagnosis, 1=one month after diagnosis, 2=3-4 months after diagnosis. EM=erythema migrans rash, NA=Not applicable. Archived sera samples were not available for all time points in each subject.
