## Supplemental Table 2 for "Secretory leukocyte protease inhibitor influences periarticular joint inflammation in *B. burgdorferi*-infected mice"

**Table S2. The primers used in this study.**

| **Gene name** | **Primer sequence** |
| --- | --- |
| Mouse *β-actin* | F: AGCGGGAAATCGTGCGTG  R: CAGGGTACATGGTGGTGCC |
| *Borrelia* *flaB* | F: TTCAATCAGGTAACGGCACA  R: GACGCRRGAGACCCTGAAAG |
| Mouse KC qPCR | F: GGCGCCTATCGCCAATG R: CTGGATGTTCTTGAGGTGAATCC |
| Mouse MCP1 qPCR | F: GTTGGCTCAGCCAGATGCA R: AGCCTACTCATTGGGATCATCTTG |
| Mouse CCR2 qPCR | F: AGTAACTGTGTGGATTGACAAGCACTTAGA R: CAACAAAGGCATAAATGACAGGAT |
| Mouse CXCR2 qPCR | F: CACCCTCTTTAAGGCCCACAT R: ACAAGGACGACAGCGAAGATG |
